# A neuronal prospect theory model in the brain reward circuitry

**DOI:** 10.1101/2021.12.18.473272

**Authors:** Yuri Imaizumi, Agnieszka Tymula, Yasuhiro Tsubo, Masayuki Matsumoto, Hiroshi Yamada

## Abstract

Prospect theory, arguably the most prominent theory of choice, is an obvious candidate for neural valuation models. How the activity of individual neurons, a possible computational unit, reflects prospect theory remains unknown. Here, we show with theoretical accuracy equivalent to that of human neuroimaging studies that single-neuron activity in four core reward-related cortical and subcortical regions represents the subjective valuation of risky gambles in monkeys. The activity of individual neurons in monkeys passively viewing a lottery reflects the desirability of probabilistic rewards, parameterized as a multiplicative combination of a utility and probability weighting functions in the prospect theory framework. The diverse patterns of valuation signals were not localized but distributed throughout most parts of the reward circuitry. A network model aggregating these signals reliably reconstructed risk preferences and subjective probability perceptions revealed by the animals’ choices. Thus, distributed neural coding explains the computation of subjective valuations under risk.

## INTRODUCTION

Prospect theory (Kahneman and Tversky, 1979) proposes that people calculate subjective valuations of risky prospects by a multiplicative combination of their subjective perceptions of two aspects of rewards: a value function that captures the desirability of rewards (i.e., utility) and an inverse S-shaped probability weighting function (i.e., probability weight) that captures a person’s subjective perception of the reward probability. Prospect theory has been the predominant model for describing human choice behavior. The nascent field of neuroeconomics has made significant progress toward an understanding of how the brain makes economic decisions (Camerer et al., 2005; Glimcher et al., 2008); however, many questions remain. One of the fundamental questions is whether discharges from individual neurons follow the prospect theory model.

Human neuroimaging provides fundamental insights into how economic decision-making is processed by brain activity, especially in the reward circuitry across cortical and subcortical structures (Haber and Knutson, 2010). This circuitry is thought to learn the values of rewards and the probability of receiving them through experience (Montague et al., 1996; Schultz et al., 1997) and it allows human decision-makers to compute subjective valuations of options. To establish a biologically viable, unified framework explaining economic decision-making, neuroeconomists have applied prospect theory to search for subjective value signals in the human brain using neuroimaging techniques (Hsu et al., 2009; Tobler et al., 2008; Tom et al., 2007). Focusing on the gain domain, previous studies found that the activity of brain regions in the reward circuitry correlates with individual subjective valuations as proposed by the prospect theory (Abler et al., 2006; Berns et al., 2008; Preuschoff et al., 2006; Tobler et al., 2008). However, limitations in temporal and spatial resolutions in neuroimaging techniques have restricted our understanding of how the reward circuitry computes subjective valuations of economic decisions, and there have been almost no studies involving the prospect theory analysis of neural mechanisms in the last decade.

Recordings of single-neuron activity in monkeys during gambling behavior may offer substantial progress over existing neuroimaging studies (Abler et al., 2006; Berns et al., 2008; Preuschoff et al., 2006; Tobler et al., 2008). Compared to human research, internal valuation measurements of probabilistic rewards have so far been limited in animals, and not all aspects of the prospect theory model could have been measured (e.g., (Yamada et al., 2013b) used only a single probability of 0.5). Recent studies have extended this earlier work asking whether captive macaques also distort probabilities in the same way humans do (Farashahi et al., 2018; Ferrari-Toniolo et al., 2019; Nioche et al., 2021; Stauffer et al., 2015), but no research has identified yet whether the activity of individual neurons in the reward circuitry computes the subjective valuation of risky prospects in a way that is consistent with prospect theory.

Thus, we targeted the reward-related cortical and subcortical structures of non-human primates (Haber and Knutson, 2010): central part of the orbitofrontal cortex (cOFC, area 13M), medial part of the orbitofrontal cortex (mOFC, area 14O), dorsal striatum (DS, the caudate nucleus), and ventral striatum (VS). We measured the neural activity in a non-choice situation while monkeys perceived a lottery with a range of probability and magnitude of rewards (10 reward magnitudes by 10 reward probabilities, resulting in 100 unique lotteries). We found neurons whose activity can be parameterized using the prospect theory model as a multiplicative combination of subjective value (utility) and subjective probability (probability weighting) functions. A simple network model that aggregates these subjective valuation signals via linear integration successfully reconstructed the monkey’s risk preference and subjective probability perception estimated from choices monkeys made in other situations. This is an evidence for a neuronal prospect theory model employing distributed computations in the reward circuitry.

## RESULTS

### Prospect theory and decision characteristics in monkeys

We estimated the monkeys’ subjective valuations of risky rewards using a gambling task (Figure 1A) (Yamada et al., 2021) similar to those used with human subjects in economics (Hey and Orme, 1994). In the choice trials, monkeys chose between two options that offered an amount of liquid reward with some probability. The monkeys fixated on a central gray target, and then, two options were presented visually as pie charts displayed on the left and right sides of the screen. The number of green pie segments indicated the magnitude of the liquid reward in 0.1 mL increments (0.1–1.0 mL), and the number of blue pie segments indicated the probability of receiving the reward in 0.1 increments (0.1–1.0, where 1.0 indicates a 100% chance). The monkeys chose between the left and right targets by fixating on one side. Following the choice, the monkeys received or did not receive the amount of liquid reward associated with their chosen option according to their corresponding probability. In each choice trial, two out of the 100 possible combinations of probability and magnitude of rewards were randomly selected and allocated to the left- and right-side target options. We used all data collected after each monkey learned to associate the probability and magnitude with the pie-chart stimuli. This included 44,883 decisions made by monkey SUN (obtained in 884 blocks over 242 days) and 19,292 decisions by monkey FU (obtained in 571 blocks over 127 days). These well-trained monkeys, like humans, showed behavior consistent with utility maximization, selecting on average options with the higher expected value, i.e., probability times magnitude (Figure 1B). In the experiment, a block of choice trials was occasionally interleaved with a block of single-cue trials (Figure 1C), during which neural recordings were made. In these trials, the monkey did not make a choice but passively viewed a single lottery cue, which offered some amount of reward with some probability given after a delay.

**Figure 1.**
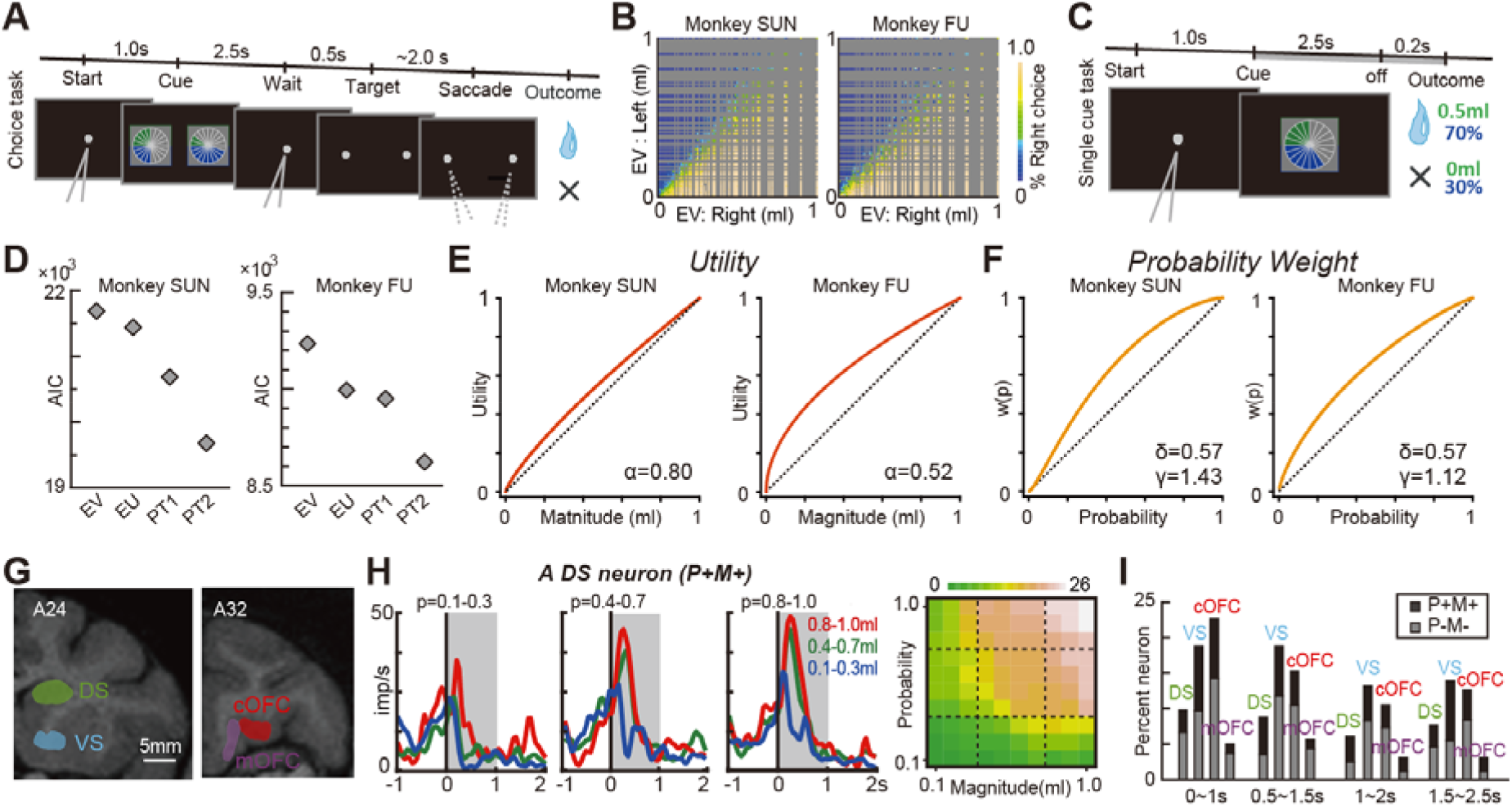
Cued lottery task, monkeys’ choice behavior, and neural coding of probability and magnitude of rewards. (A) A sequence of events in the choice trials. Two pie charts representing the available options were presented to the monkeys on the left and right sides of the screen. Monkeys chose either of the targets by fixating on the side where it appeared. (B) The frequency with which the target on the right side was selected for the expected values of the left and right target options. (C) A sequence of events in the single-cue trials. (D) AIC values are estimated based on the four standard economic models to describe monkey’s choice behavior: EV, EU, PT1, and PT2. See Methods for details. (E) Estimated utility functions in the best-fit model PT2. (F) Estimated probability weighting functions in the best-fit model PT2. (G) An illustration of neural recording areas based on coronal magnetic resonance images. (H) Example activity histogram of a DS neuron modulated by the probability and magnitude of rewards with positive regression coefficients during the single-cue task. The activity aligned to the cue onset is represented for three different levels of probability (0.1– 0.3, 0.4–0.7, 0.8–1.0) and magnitude (0.1–0.3 mL, 0.4–0.7 mL, 0.8–1.0 mL) of rewards. Gray hatched time windows indicate the 1-s time window used to estimate the neural firing rates shown in the right graph displaying the average smoothing between neighboring pixels. (I) Percentage of neurons modulated by probability and magnitude of rewards in the four core reward brain regions. Black indicates activity showing positive regression coefficients for probability and magnitude of rewards (P+M+ type). Gray indicates activity showing the negative regression coefficients for probability and magnitude (P-M- type).(A)–(C) and (G) have been previously published in Yamada et al., 2021.

We estimated each monkey’s utility and probability weighting functions from their choice behavior using the standard parametrizations in the literature. For the utility function, we used the power utility function *u(m) = m*^*α*^, where *m* indicates the magnitude of reward, *α* > 1 indicates convex utility (risk-seeking behavior), *α* < 1 indicates concave utility (risk aversion), and *α* = 1 indicates linear utility (risk neutrality). For the probability weighting function *w(p)*, we used one-parameter, *w(p) = exp(- (-log p)^γ^)*, and two-parameter, *w(p) = exp(-δ (-log p)^γ^)*, Prelec functions. The one-parameter version is nested in the two-parameter version (when *δ* = 1) for ease of comparison. Overall, we estimated the following four models of the utility of receiving reward magnitude *m* with probability *p, V(p,m)*:

1. EV: expected value *V(p,m) = p m*
2. EU: expected utility *V(p,m) = p m*^*α*^
3. PT1, one-parameter Prelec: prospect theory with *w(p)* as in (Wu and Gonzalez, 1996)

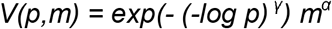
4. PT2, two-parameter Prelec: prospect theory with *w(p)* as in (Prelec, 1998)

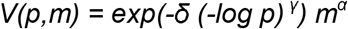

*α, δ*, and *γ* are free parameters, and *p* and *m* are the probability and magnitude of reward cued by the lottery, respectively. The parameters *δ* and *γ* control the subproportionality and regressiveness of *w(p)*. We assumed that subjective probabilities and utilities are integrated multiplicatively, as is customary in economic theory, yielding *V(p,m) = w(p) u(m)*. The probability of the monkey choosing the lottery on the right side (*L*_*R*_) instead of the lottery on the left side (*L*_*L*_) was estimated using a logistic choice function:

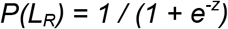

where *z = β (V(L*_*R*_*) - V(L*_*L*_)), and the free parameter *β* controls the degree of stochasticity observed in the choices.

To determine which model best describes the behavior of a monkey, we used Akaike’s Information Criterion (AIC), which measures the goodness of model fit with a penalty for the number of free parameters employed by the model (see Methods for more details). Among the four models, PT2 had the lowest AIC and outperformed EV, EU, and PT1 in both monkeys (Figure 1D). In the best-fit model, the utility function was concave (Figure 1E; one-sample t-test, *α* = 0.80, z = 46.10, P < 0.001 in monkey SUN; *α* = 0.52, P < 0.001, z = 25.04 in monkey FU), indicating that monkeys were risk-averse. Notably, for both monkeys, the probability weighting functions were concave instead of the inverse-S shape traditionally assumed in humans (Figure 1F; one-sample t-test, *δ* = 0.57, z = 86.51, P < 0.001 in monkey SUN; *δ* = 0.57, z = 52.77, P < 0.001 in monkey FU; *γ* = 1.43, z = 47.29, P < 0.001 in monkey SUN; *γ* = 1.12, z = 25.68 in monkey FU, P < 0.001). Overall, we conclude that in monkeys, utility functions estimated from behavior are concave, similar to those in humans, but monkeys distort probability differently compared to what is usually assumed for human decision-makers.

### Neural signals for subjective valuations are distributed in the reward circuitry

We recorded single-neuron activity during the single-cue task (Figure 1C) from neurons in the DS (n=194), VS (144), cOFC (190), and mOFC (158) (Figure 1G). These brain regions are known to be involved in decision-making. We first identified neurons whose activity represents the key reward statistics – probability and magnitude – that underlie the expected value, expected utility, and prospect theory. These neurons were identified by regressing neural activity on probability and magnitude of rewards, and the neurons included in our analysis were those that had either both positive or both negative regression coefficients (See Methods).

An example of activity during a one-second time window after cue onset is shown in Figure 1H. This DS neuron showed an activity modulated by both the probability and magnitude of rewards with positive regression coefficients (P+M+ type; probability, regression coefficient, r = 13.51, t = 8.57, P < 0.001; magnitude, r = 12.27, t = 7.79, P < 0.001). Neuronal firing rates increased as the reward probability increased and as the reward magnitude increased, representing a positive coding type (Figure 1H, right). Similarly, some neurons showed an activity modulated by both the probability and magnitude of rewards with negative regression coefficients, representing a negative coding type (P-M- type). In total, these types of activity were observed in 24% (164/686) of all recorded neurons in at least one of the four analysis epochs during the 2.5-s cue period. The proportions of these signals in each brain region were different (DS, 22%, 43/194, VS, 32%, 45/141, cOFC, 31%, 59/190, mOFC, 11%, 17/158, chi-squared test, Χ^2^ = 25.59, df = 3, P < 0.001). These neurons were evident across the entire cue period (Figure 1I), during which the monkeys perceived the probability and magnitude of rewards.

### Detecting the neuronal signature of prospect theory

For visual inspection of the potential neuronal signature of *V(p,m)*, we predicted from the behavioral estimates how the observed neuronal firing rates should look like in each of the four models: expected value (Figure 2A, EV), expected utility (Figure 2B, EU), and prospect theory (Figure 2C and 2D, PT1 and PT2, respectively). In each of the models, the neural firing rate *R* is given by:

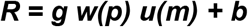

where the predicted neuronal responses *R*, the output of the model, integrates the subjective value function (i.e., utility, *u(m)*) and subjective probability function (i.e., probability weight, *w(p)*). *b* is a free parameter that captures the baseline firing rates in the probability-magnitude space. *g* determines the magnitude of the neural responses to *u(m)* and *w(p). u(m)* and *w(p)* are specified for each model as described above (see the formulas in Figure 2 and Methods).

**Figure 2.**
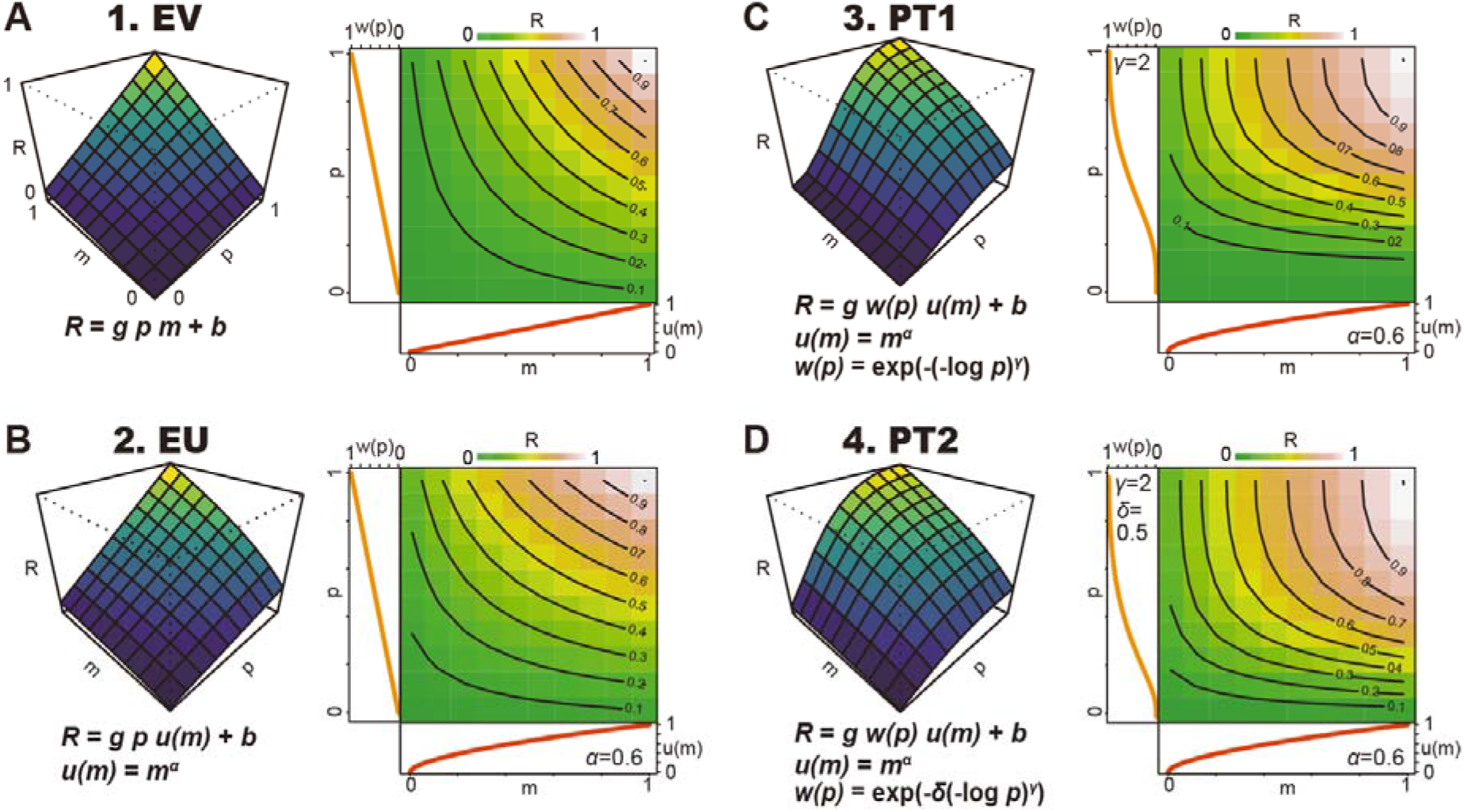
Neural models of economic decision theory. Schematic depiction of predicted neuronal responses *R* defined by the four economic models that represent expected value (A, EV), expected utility (B, EU), and prospect theory one-parameter Prelec (C, PT1) and two-parameter Prelec (D, PT2). Model equations are shown in each plot. *R* is plotted against the probability (*p*) and magnitude (*m*) of the rewards. *b, g, α, γ*, and *δ* are free parameters. *g* and *b* are the gain and intercept parameters, respectively. *α* represents the curvature of the *u(m)*. *δ* and *γ* represent probability weighting functions. For these schematic drawings, the following values for free parameters were used: *b, g, α, γ*, and *δ* were 0 spk s^-1^, 1, 0.6, 2, and 0.5, respectively, for all four figures. See Methods section for more details.

Next, we aimed to assess which of the models best captures the neuronal discharge rates in each brain region. Therefore, we fitted the activities of individual neurons with each of the four models, treating *b, g, *α*, δ*, and *γ* as free parameters. Our carefully designed set of lottery stimuli – a sampling matrix of 10 rewards by 10 probabilities – allowed us to perform a reliable estimation of these five free parameters for each activity of neurons. To determine which model best describes the observed neuronal firing rate in each individual neurons, we used the AIC. As demonstrated for an example neuron in Figure 3A, the activity of this DS neuron was best explained by prospect theory with a two-parameter probability weighting function (Figure 3B, PT2). For this neuron, PT2 had the smallest AIC values with the highest percentage of explained variance. The output *R* of the fitted PT2 model described the activity pattern well (Figure 3C), as well as the observed activity (Figure 3A), in which the neural utility function and subjective probability weighting function were parameterized (Figure 3D) via a multiplicative relation in the model.

**Figure 3.**
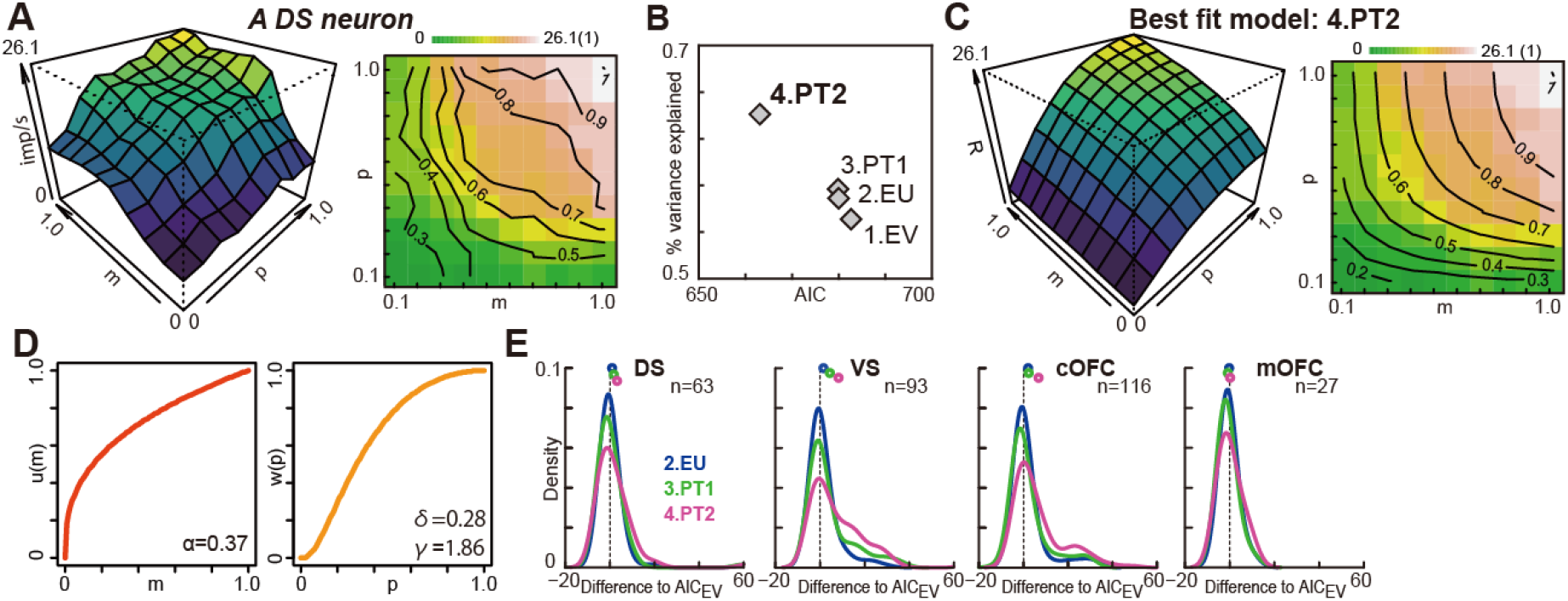
Prospect theory best explained neural firing rates in the reward circuitry. (A) Plot of an example activity of the DS neuron in Figure 1H against probability (*p*) and magnitude (*m*) of rewards. To draw the 3D curvature (left) and contour lines (right), neighbouring pixels were average smoothed. (B) The AIC values against the percent variance explained are plotted in each model for the example neuron in (A). (C) A 3D histogram (left) and contour lines (right) predicted from the best-fit PT2 model in (A). The activity of the example neuron in (A) is shown in the right color map figure. Contour lines are shown for every 10% change in the fitted model. (D) *u(m)* and *w(p)* estimated in the best-fit model PT2 for the neural activity in (A). (E) Probability density of the estimated AIC difference of the three models against the EV (the simplest) model. The plots display mean values. n represents the number of neuronal signals that showed both positive or both negative regression coefficients for probability and magnitude of rewards.

To understand which model best describes the neural activity in each brain region, we determined the goodness-of-fit score for each activity of the neurons as the difference in AIC between each of the models (EU, PT1, and PT2) and the EV model. Here, we treated the EV model as the baseline because it is the simplest model and a predecessor of the other models in the economics literature. Figure 3E shows the probability density of the goodness-of-fit score differences for each brain region separately. The vertical dashed lines at zero indicate no difference in the AIC of the EV model and that of the model under consideration. A model that shows more deviation to the right of the graph indicates a better fit.

Overall, prospect theory (PT2) best described the activity of most neural populations in the reward circuitry (DS, VS, and cOFC), except for mOFC activity. We statistically compared the AIC values among the four models. The comparisons indicated that the PT2 model was best at describing DS, VS, and cOFC activity as a whole (one-sample t-test after subtracting models’ AIC scores; DS: df = 62, EV-EU, t = 0.94, P = 0.35, EU-PT1, t = 1.03, P = 0.31, PT1-PT2, t = 3.01, P = 0.004; VS: df = 92, EV-EU, t = 2.42, P = 0.017, EU-PT1, t = 4.00, P < 0.001, PT1-PT2, t = 3.91, P < 0.001; cOFC: df = 115, EV-EU, t = 2.90, P = 0.004, EU-PT1, t = 0.65, P = 0.52, PT1-PT2, t = 6.18, P < 0.001, not shown for all). However, the best descriptive model of the mOFC activity could not be determined (one-sample t-test; mOFC: df = 26, EV-EU, P = 0.60, EU-PT1, P = 0.10, PT1-PT2, P = 0.11), suggesting that mOFC neurons simply signal expected values, without any distortions to objective probability and magnitude of rewards during the perception of the lottery.

Next, we asked whether neurons differentially encode subjective valuations based on their location (DS, VS, and cOFC). For this purpose, we used the PT2 model estimates *b, g, α*, δ, and *γ* of individual activity of neurons, including both positive and negative coding types. We clustered these five parameters using k-means clustering algorithms following principal component analysis (PCA) across the neural population in the DS, VS, and cOFC (Figure 4A and 4B, see Methods). The five predominant clusters, C1 to C5, were obtained after PCA based on the four principal components (Figure 4B). These five clusters were observed in similar proportion across the three brain regions with only slight differences (Figure 4C). One small difference was that the VS contained a smaller proportion of the predominant cluster than the other two regions (chi-squared test, Χ^2^ = 18.15, df = 8, P = 0.020).

**Figure 4.**
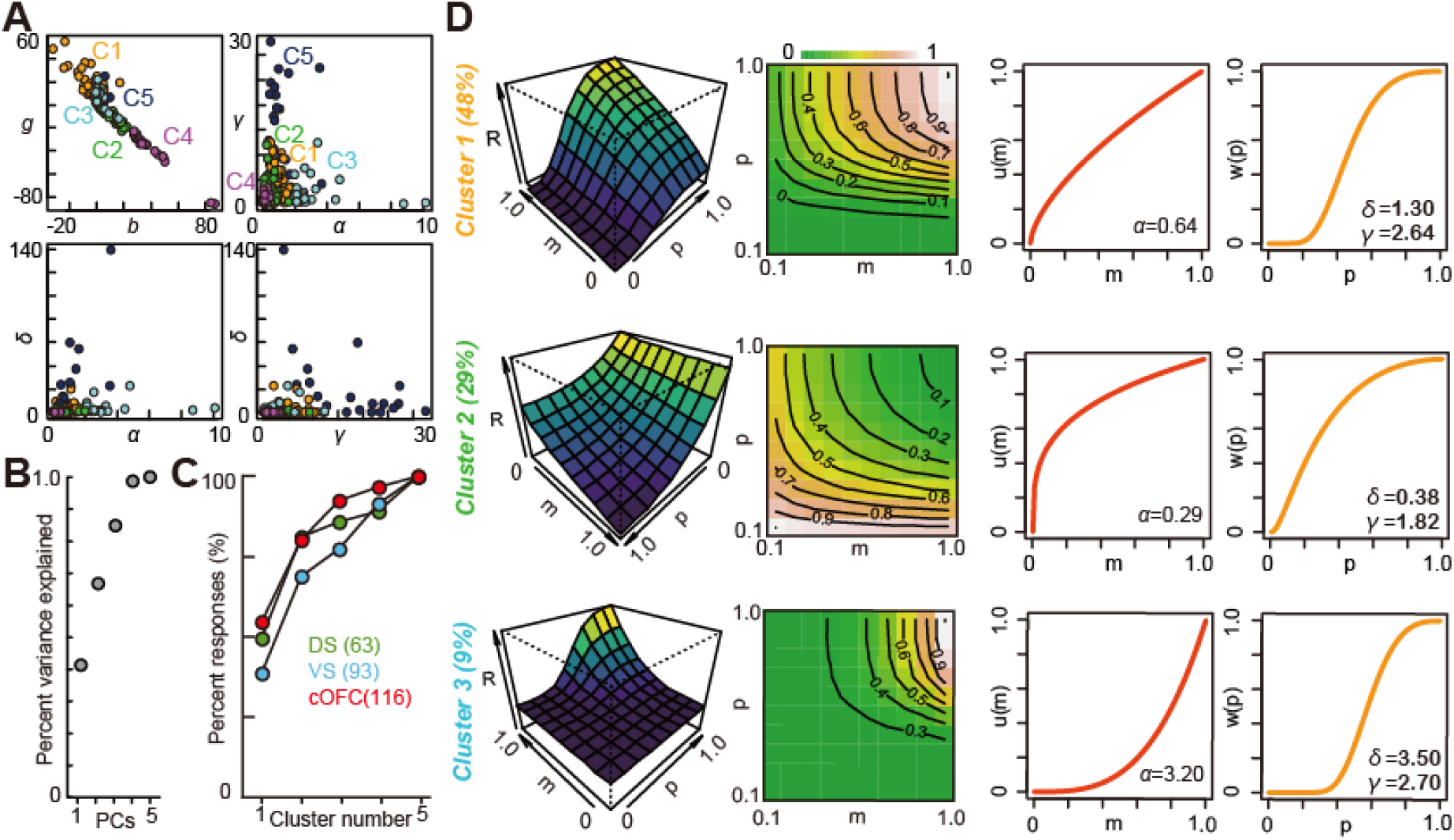
Neuronal clusters categorized by the fitted parameters according to the prospect theory model. (A) Plots of all five parameters estimated in DS, VS, and cOFC neurons. *g, b, α*, δ, and *γ* are plotted. (B) Cumulative plot of the percent variance explained by PCA is shown against the principal components PC1 to PC5. (C) Cumulative plot of the percentages of activity categorized into the five clusters in each brain region. (D) Response *R* (model output) in the first three predominant clusters are plotted. 3D curvature, contour lines with color maps, *u(m)*, and *w(p)* are plotted using mean values of each parameter in each cluster. For drawing the 3D curvature (first column) and contour lines (second column), *R* is normalized by the maximal value.

Across the DS, VS, and cOFC, the predominant cluster, C1, represented 48% of all activity (Figure 4D, top row; mean values: *b* = -0.68, *g* = 10.1, *α* = 0.64, *δ* = 1.30, *γ* = 2.64). Its output, *R*, was described by a combination of a concave utility function and an S-shaped probability weighting function (Figure 4D, see the third and fourth columns in the top row). The second predominant cluster, C2, was also best described with a concave utility function, but its probability weighting function was concave. This cluster was mostly composed of neurons with negative coding of probability and magnitude of rewards (Figure 4D, middle row; *b* = 10.6, *g* = -10.1, *α* = 0.29, *δ* = 0.38, *γ* = 1.82). Because the coding gain was negative (Figure 4D, middle left, note that axis values are plotted from 1.0 to 0), the convex curvature (Figure 4D, left column in the middle row) of the firing rate corresponds to the concave functions *u(m)* and *p(w)*. A considerable proportion of neurons (9%), C3, showed output well described by a convex utility function and an S-shaped probability weighting function with a smaller gain compared to C1 and C2 (Figure 4D, bottom; *b* = 2.6, *g* = 7.2, *α* = 3.2, *δ* = 3.5, *γ* = 2.7).

These clusters of neurons parameterized by the prospect theory model were not localized and were instead found scattered across most parts of the reward circuitry (DS, VS, and cOFC), suggesting that distributed coding underlies internal subjective valuations under risk.

### Reconstruction of internal preference parameters from observed neural activity in monkeys

Lastly, we reconstructed the monkeys’ internal valuations of passively viewed lotteries from the observed neural activity to assess how well they match the utility and probability weighting functions estimated from the behavioral choices. To do so, we constructed a simple three-layered network model as a minimal rate model, a primitive version of the advanced models used recently (Juslin et al., 2003; Ohshiro et al., 2011), and simulated the choices of this network model (Figure 5). We assumed that outputs reflecting *V(p,m)* in each neural cluster C1 to C5 (Figure 5A, first layer) were linearly integrated by the network (Figure 5A, second layer, population SEVs, see Methods). The activities in clusters 1, 3, and 5 (mostly composed of P+M+ neurons) were linearly summed, and the activities in clusters 2 and 4 (mostly composed of P-M-neurons) were subtracted to integrate the opposed signals (hence, linear summation of an inversed signal). To simulate choice, we generated two identical population SEVs for the left (ΣR_L_) and right (ΣR_R_) target options and used a random utility model for selecting one option (Figure 5A, third layer, sigmoid choice function). Overall, we simulated 40,000 choices – four times each possible combination of 100 lotteries, *L(p,m)*.

**Figure 5.**
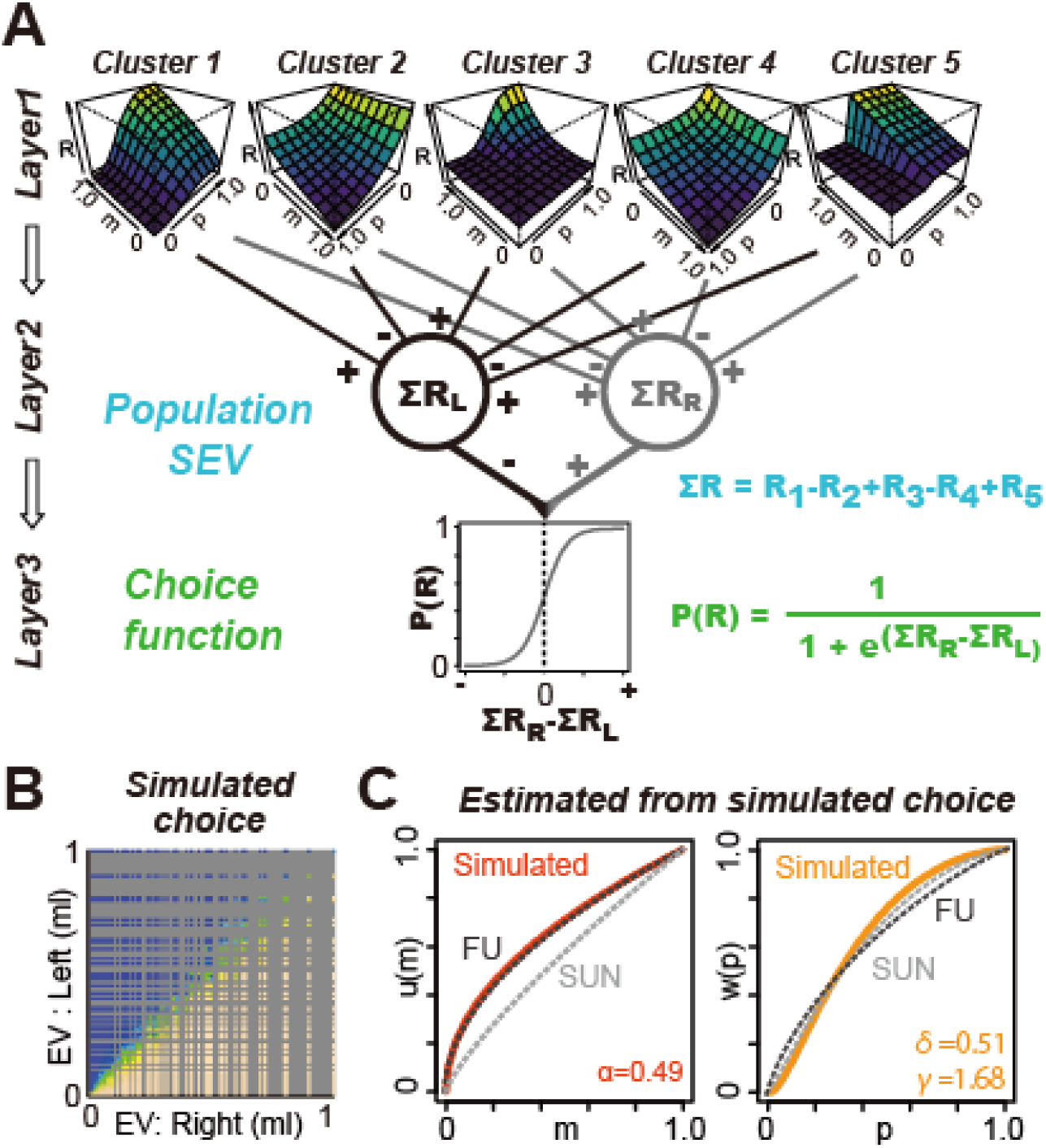
A simple network model reconstructs the subjective decision statistics in monkeys. (A) The five neural clusters as detected by PCA in the reward circuitry. Subjective expected value functions (SEVs) for left and right target options are defined as the linear summation of the five clusters (see Methods). Choice is simulated as a sigmoid function of the subjective value signal difference. (B) The frequency with which the target on the right side was selected by a computer simulation based on the network shown in (A). (C) *u(m)* and *w(p)* estimated from the simulated choice in (B) are plotted. Dotted lines indicate the actual functions *u(m)* and *w(p)* of the monkeys, as shown in Figure 1E and 1F, respectively.

While our network model used neural signals modeled by prospect theory during passive viewing, these simulated choice patterns based on the clustered neuronal prospect theory model were very similar to the actual gambling behaviors of the monkeys (Figures 5B and 1B). When estimating the utility function and probability weighting function of these simulated choices, we observed concave utility functions and concave probability weighting functions similar to those obtained from the actual gambling behavior (Figure 5C). Thus, we conclude that a distributed neural code that accumulates individual neuronal signals can explain the internal subjective valuations of monkeys.

## Discussion

Prospect theory is the dominant theory of choice in behavioral economics, but it remains elusive whether the theory is only descriptive of human behavior or has a deeper meaning in the sense that it also describes an underlying neuronal computation that extends to other species. Previous human neuroimaging studies have demonstrated that neural responses to rewards measured through blood oxygen levels can be described using prospect theory (Hsu et al., 2009; Tobler et al., 2008; Tom et al., 2007) but with limited resolution in temporal and spatial domains. Here, we provided the first evidence that the activity of individual neurons in the reward circuitry (DS, VS, and cOFC) of monkeys perceiving a lottery can be captured based on the prospect theory model as a multiplicative combination of utility and probability weighting functions (Figure 4). One pivotal question is how these various subjective preference signals are transformed into behavioral choices through information processing via neural networks. Our clustering analysis of the parameterized neuronal activity revealed that these signals were similarly distributed across the VS, DS, and cOFC (Figure 4C). Our minimal rate model of a three-layered network successfully reconstructed the internal valuation of risky rewards observed in monkeys (Figure 5), suggesting that these subjective valuation signals in the reward circuitry are integrated into the brain to construct a decision output from risky perspectives.

Previous studies have shown that neuronal signals related to cognitive and motor functions are widely observed in many brain regions (Bouton et al., 2018; Coghill, 2020; Nestor et al., 2011; Pinel et al., 2004; Simon et al., 2006; Stefanini et al., 2020; Wixted et al., 2014). These distributed neuronal signals suggest that a distributed neural code is a common computation in the brain. The recent development of large-scale neural recording technologies verified that this is a common computational mode (Steinmetz et al., 2019); the analysis of approximately 30,000 neurons in 42 regions of the rodent brain revealed that behaviorally relevant task parameters are observed throughout the brain. Our results from the reward-related brain regions are in line with this view, except for the mOFC, where fewer encodings of probability and magnitude of rewards were observed (Figures 1I and 3E). This might be because the medial-lateral axis in the reward circuitry yields a significant difference in reward-based decision-making (Haber and Knutson, 2010). The distributed code may require some input-output functions (Vankov and Bowers, 2017) to process the probability and magnitude of rewards and integrate these information to estimate the expected subjective utility, at least in some neural populations. One possible information processing for this input-output mapping can be achieved by neural population dynamics (Chen and Stuphorn, 2015; Gardner et al., 2019; Yoo and Hayden, 2020), in which some subclusters of neurons can process information moment-by-moment as a dynamical system. Stable neural population dynamics in the VS and cOFC were indeed observed in contrast to the fluctuating signals in the DS population (Yamada et al., 2021), which may reflect some differences in distributed coding.

One limitation of our study is that our application of prospect theory is limited to the domain of gains, since unlike in human studies that use money as the reward, it is impossible to take fluid rewards from monkeys to make them experience losses. Nevertheless, our study adds important behavioral evidence to the growing literature on prospect theory preferences in primates. Recent studies of captive macaques have begun to investigate distortions in the perception of probabilities, with inconsistent results across studies (Eisenreich et al., 2019; Farashahi et al., 2018; Ferrari-Toniolo et al., 2019; Nioche et al., 2019; Nioche et al., 2021; Stauffer et al., 2015). The probability weighting function was inverse S-shaped (Farashahi et al., 2018; Ferrari-Toniolo et al., 2019), S-shaped (Nioche et al., 2021; Stauffer et al., 2015), or concave (Ferrari-Toniolo et al., 2021; Ferrari-Toniolo et al., 2019). Although we consistently found that the probability weighting functions of our two well-trained monkeys were concave, most studies conducted in humans have found inverse-S-shaped probability weighting functions at the aggregate level, with a large amount of heterogeneity at the individual level (Abdellaoui, 2000; Bruhin et al., 2010; Fehr-Duda et al., 2011; Harbaugh et al., 2002; Harrison and Rutstrom, 2009; Hsu et al., 2009; Tobler et al., 2008) indicating an inconsistency across the two species. Furthermore, the monkeys in the present study had concave utility functions while most previous studies have found that monkeys have a convex (Farashahi et al., 2018; Stauffer et al., 2015) or concave (Eisenreich et al., 2019; Ferrari-Toniolo et al., 2021; Nioche et al., 2019; Yamada et al., 2013b) utility over rewards in the gain domain. In conclusion, our monkeys had concave utility functions, similar to our previous findings in monkeys (Yamada et al., 2018; Yamada et al., 2013b) as well as in humans. But unlike humans, our monkeys had concave probability weighting functions.

Summing up, we provided novel evidence that the activity of the individual neurons in the reward circuitry can be described using prospect theory and that the probability distortions estimated from the monkeys’ behaviors are different than those usually assumed for humans.

## METHODS

### Subjects and experimental procedures

Two rhesus monkeys performed the task (*Macaca mulatta*, SUN, 7.1 kg, male; *Macaca fuscata*, FU, 6.7 kg, female). All experimental procedures were approved by the Animal Care and Use Committee of the University of Tsukuba (Protocol No H30.336) and performed in compliance with the US Public Health Service’s Guide for the Care and Use of Laboratory Animals. Each animal was implanted with a head-restraint prosthesis. Eye movements were measured using a video camera at 120 Hz. Visual stimuli were generated by a liquid-crystal display at 60 Hz, placed 38 cm from the monkey’s face when seated. The subjects performed the cued lottery task five days a week. The subjects practiced the cued lottery task for 10 months, after which they became proficient in choosing lottery options.

### Cued lottery tasks

Animals performed one of two visually cued lottery tasks: a single-cue task or a choice task.

#### Single-cue task

At the beginning of each trial, the monkeys had 2 s to align their gaze within 3° to a 1°-diameter gray central fixation target. After fixing for 1 s, an 8° pie chart providing information about the probability and magnitude of rewards was presented for 2.5 s at the same location as the central fixation target. Probability and magnitude were indicated by the numbers of blue and green pie chart segments, respectively. The pie chart was then removed and 0.2 s later, a 1-kHz and 0.1-kHz tone of 0.15-s duration indicated reward and no-reward outcomes, respectively. The high tone preceded reward delivery by 0.2 s, whereas the low tone indicated that no reward was delivered. The animals received a liquid reward, as indicated by the number of green pie chart segments with the probability indicated by the number of blue pie chart segments. An intertrial interval of 4–6 s followed each trial.

#### Choice task

At the beginning of each trial, the monkeys had 2 s to align their gaze within 3° to a 1°-diameter gray central fixation target. After fixation for 1 s, two peripheral 8° pie charts providing information about the probability and magnitude of rewards for each of the two target options were presented for 2.5 s at 8° to the left and right of the central fixation location. The gray 1° chosen targets appeared at the same locations. After a 0.5-s delay, the fixation target disappeared, cueing saccade initiation. The monkeys were allowed 2 s to make their choice by shifting their gaze to either target within 3° of the chosen target. A 1-kHz and 0.1-kHz tone sounded for 0.15 s to denote reward and no-reward outcomes, respectively. The animals received a liquid reward, as indicated by the number of green pie chart segments of the chosen target with the probability indicated by the number of blue pie chart segments. An intertrial interval of 4–6 s followed each trial.

#### Payoff, block structure, and data collection

Green and blue pie charts respectively indicated reward magnitudes from 0.1 to 1.0 mL, in 0.1 mL increments, and reward probabilities from 0.1 to 1.0, in 0.1 increments. A total of 100 pie chart combinations were used. In the single-cue task, each pie chart was presented once in a random order, allowing monkeys to experience all 100 lotteries within a certain period. In the choice task, two pie charts were randomly allocated to the left and right targets in each trial. Approximately 30–60 trial blocks of the choice task were sometimes interleaved with 100–120 trial blocks of the single-cue task.

#### Calibration of the reward supply system

A precise amount of liquid reward was delivered to the monkeys using a solenoid valve. An 18-gauge tube (0.9 mm inner diameter) was attached to the tip of the delivery tube to reduce the variation across trials. The amount of reward in each payoff condition was calibrated by measuring the weight of water with 0.002 g precision (2 μL) on a single-trial basis. This calibration method was the same as that used in (Yamada et al., 2018).

#### Electrophysiological recordings

We used conventional techniques to record single-neuron activity from the DS, VS, cOFC, and mOFC. Monkeys were implanted with recording chambers (28 × 32 mm) targeting the OFC and striatum, centered 28 mm anterior to the stereotaxic coordinates. The locations of the chambers were verified using anatomical magnetic resonance imaging. We used a tungsten microelectrode (1–3 MΩ, FHC) to record the neurons. Electrophysiological signals were amplified, band-pass-filtered, and monitored. Single-neuron activity was isolated based on the spike waveforms. We recorded from the four brain regions of a single hemisphere of each of the two monkeys: 194 DS neurons (98 and 96 from monkeys SUN and FU, respectively), 144 VS neurons (89, SUN and 55, FU), 190 cOFC neurons (98, SUN and 92, FU), and 158 mOFC neurons (64, SUN and 94, FU). The activity of all single neurons was sampled when the activity of an isolated neuron demonstrated a good signal-to-noise ratio (> 2.5). Blinding was not performed. The sample sizes required to detect effect sizes (the number of recorded neurons, the number of recorded trials in a single neuron, and the number of monkeys) were estimated in reference to previous studies (Chen and Stuphorn, 2015; Yamada et al., 2013a; Yamada et al., 2018). Neural activity was recorded during 100–120 trials of the single-cue task. During the choice trials, the neural activity was not recorded. Presumed projection neurons (phasically active neurons, (Yamada et al., 2016)) were recorded from the DS and VS, whereas presumed cholinergic interneurons (tonically active neurons, (Inokawa et al., 2020; Yamada et al., 2004) were not recorded.

### Statistical analysis

For statistical analysis, we used the statistical software R and Stata. All statistical tests were two-tailed. We used standard maximum likelihood procedures to estimate utility functions and probability weighting functions in Stata. We performed a neural analysis and simulation to reconstruct the choice from a neural model in R.

### Behavioral analysis

We first examined whether the choice behavior of a monkey depended on the expected values of the two options located on the left and right sides of the screen. We pooled choice data across all recording sessions (monkey SUN, 884 sessions, 242 days; monkey FU, 571 sessions, 127 days), yielding 44,883 and 19,292 choice trials for monkeys SUN and FU, respectively. The percentage of the right target choices was estimated from the pooled choice data for all combinations of the expected values of the left and right target options. This result has been reported previously (Yamada et al., 2021).

### Economic models

We estimated the parameters of the utility and probability weighting functions within a random utility framework. Specifically, a lottery *L(p,m)* denoted a gamble that pays *m* (magnitude of the offered reward in mL) with a probability *p* or 0 otherwise. We assumed a popular constant relative risk attitude (CRRA, also known as power utility function), *u(m) = m*^*α*^, and considered the previously proposed probability weighting functions. We assumed two subjective probability functions *w(p)* commonly used in the prospect theory; one-parameter Prelec (PT1): *w(p) = exp(- (-log p)^γ^)* (Wu and Gonzalez, 1996) and two-parameter Prelec (PT2): *w(p) = exp(-*δ *(-log p)^γ^)* (Prelec, 1998). We assumed that subjective probabilities and utilities are integrated multiplicatively per standard economic theory, yielding the expected subjective utility function *V(p,m) = w(p) u(m)*.

The probability of a monkey choosing the lottery on the right side (L_R_) instead of the lottery on the left side (L_L_) was estimated using a logistic choice function:

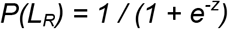

where *z = β (V(L*_*R*_*) - V(L*_*L*_)), and the free parameter *β* controls the degree of stochasticity observed in the choices. We fitted the data by maximizing log-likelihood and choosing the best structural model to describe the monkeys’ behavior using the AIC (Burnham and Anderson, 2004).

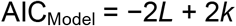

where *L* is the maximum log-likelihood of the model, and *k* is the number of free parameters.

In each fitted model, whether *α*, δ, and *γ* were significantly different from zero was determined using a one-sample t-test at P < 0.05. Whether *α*, δ, and *γ* were significantly different from one was also determined using a one-sample t-test at P < 0.05.

### Neural analysis

#### Basic firing properties

Peristimulus time histograms were drawn for each single-neuron activity aligned at the onset of a visual cue. The average activity curves were smoothed using a 50-ms Gaussian kernel (σ = 50 ms). Basic firing properties, such as peak firing rates, peak latency, and duration of peak activity (half peak width), were compared among the four brain regions using parametric or nonparametric tests, with a statistical significance level of P < 0.05. Baseline firing rates during 1 s before the appearance of central fixation targets were also compared with a statistical significance level of P < 0.05. These basic firing properties have been described in Yamada et al., 2021.

We analyzed neural activity during a 2.5-s period during pie chart stimulus presentation in the single-cue task. We estimated the firing rates of each neuron during the 1-s time window every 0.5 s after the onset of the cue stimuli. No Gaussian kernel was used.

#### Pre-screening neural activity for economic model fits

To determine which neurons were sensitive to the probability and magnitude cued by a lottery, without assuming any specific model, neural discharge rates (*F*) were regressed on a linear combination of a constant and the probability and magnitude of rewards:

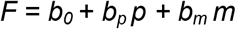

where *p* and *m* are the probability and magnitude of the rewards indicated by the pie chart, respectively. *b*_0_ is the intercept. If *b*_*p*_ and *b*_*m*_ were not 0 at P < 0.05, the discharge rates were regarded as significantly modulated by that variable.

Based on the linear regression, two types of neural modulations were identified: the “P+M+” type with a significant *b*_p_ and a significant *b*_m_ both having a positive sign (i.e., positive *b*_p_ and positive *b*_m_) and the “P-M-” type with a significant *b*_p_ and a significant *b*_m_ both having a negative sign (i.e., negative *b*_p_ and negative *b*_m_). Both types of the neuronal signal could represent the economic decision statistics described in the next section.

#### Neural economic models

We fitted the four neural models of subjective valuation of lottery *L(p,m)* to the activity of the pre-selected neurons that were sensitive to the information of probability and magnitude of rewards. The unified formula for all models is *R* = *g w(p) u(m)* + *b*, where output of the model *R* represents firing rates as a function of the subjective probability *w(p)* times the utility of reward *u(m)*, which is the subjective expected value (SEV) of a lottery that reflects the monkey’s lottery valuation. For neural representation of *V(p,m)* as described in the main text, we call this value function to differ from behavioral measures. In all models, *g* (magnitude of the neural response), *b* (baseline firing rate), *α* (utility curvature), *γ*, and *δ* (probability weighting) are free parameters.

1. ***Expected value model (EV)***.

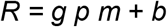
2. ***Expected utility model (EU)***.

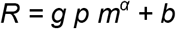
3. ***Prospect theory model with one-parameter Prelec* (PT1)**.

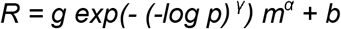
4. ***Prospect theory model with two-parameter Prelec (PT2)***.

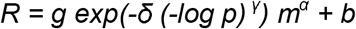

To identify the structural models that best describe the activity of neurons in each brain region, we fitted each of the models to the P+M+ and P-M-type activity of each neuron on a trial-by-trial basis. We estimated the combination of best-fit parameters using the R statistical software package. We used the nls() function in R with random initial values (repeated 100 times) to find a set of parameters that minimizes nonlinear least squared values.

For each of the four brain regions, the best-fit model showing minimal AIC was selected by comparing the AIC values among the models. If the differences in AIC values against the three other models were significantly different from zero in the one-sample t-test at P < 0.05, the model was defined as the best model. For visual presentation, we plotted AIC differences in comparison to the EV model as the baseline model in the economics literature.

#### Construction of the neural prospect theory model

The estimated parameters in the best-fit model of the neuronal activity were classified using PCA followed by the k-means clustering algorithm. PCA was applied once to all parameters estimated in the best-fit model PT2, i.e., *g, b, α, γ*, and *δ* in DS, VS, and cOFC. The k-means algorithm was used to classify five types of neural responses according to the PC1 to PC4 scores since the first four PCs explained more than 90% of the variance. Following the classification, we define each type of cluster with the mean of each estimated parameter, as the five clusters were observed in each of the DS, VS, and cOFC neural populations.

#### Evaluation of neural model performance using simulated data

We constructed a simple layered network model for simulations (Juslin et al., 2003; Ohshiro et al., 2011). We simply reconstructed a neural prospect theory model from the clusters above by adding each response *R* of the five clusters. For clusters 1, 3, and 5, we linearly summed them, while for clusters 2 and 4, which were mostly composed of P-M-types, we inversed their activity by subtraction. This population SEV was filtered by a ReLU (Rectified Linear Unit) function, since it mimics the firing rate. The linear sum of the five clusters was allocated to the left and right target options to perform a simulation based on the difference of these integrated responses. We then simulated the choice for lotteries consisting of four times of all possible combinations of lotteries *L(p,m)* using the logistic function

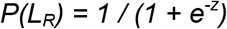

where *z = β (V(L*_*R*_*) - V(L*_*L*_)) and *β* is assumed to be one, i.e., no beta term. These simulated choice data, composed of 40,000 choice trials, were visualized and evaluated by applying the best-fit model to estimate the preference parameters *α, γ*, and *δ* in *u(m) = m*^*α*^ and *w(p) = exp(-δ (-log p)^γ^)*, as well as *β* in the choice function, similar to the model fit to the actual behavior of the monkey.

## Acknowledgments

The authors would like to thank Takashi Kawai, Ryo Tajiri, Yoshiko Yabana, and Yuki Suwa for their technical assistance, and Jun Kunimatsu and Masafumi Nejime for their valuable comments. Monkey FU was provided by the NBRP “Japanese Monkeys” through the National Bio Resource Project of the MEXT, Japan. Funding: This research was supported by JSPS KAKENHI (Grant Numbers JP:15H05374, 19H05007, and 21H02797), Takeda Science Foundation, Narishige Neuroscience Research Foundation, Research Foundation for the Electrotechnology of Chubu (H.Y.), JSPS KAKENHI 19K12165 (Y.T.), and ARC DP190100489 (A.T.).

